# Nuclear receptor interdomain communication is mediated by the hinge with ligand specificity

**DOI:** 10.1101/2024.02.10.579785

**Authors:** Saurov Hazarika, Tracy Yu, Arumay Biswas, Namita Dube, Priscilla Villalona, C. Denise Okafor

## Abstract

Nuclear receptors are ligand-induced transcription factors that bind directly to target genes and regulate their expression. Ligand binding initiates conformational changes that propagate to other domains, allosterically regulating their activity. The nature of this interdomain communication in nuclear receptors is poorly understood, largely owing to the difficulty of experimentally characterizing full-length structures. We have applied computational modeling approaches to describe and study the structure of the full length farnesoid X receptor (FXR), approximated by the DNA binding domain (DBD) and ligand binding domain (LBD) connected by the flexible hinge region. Using extended molecular dynamics simulations (> 10 microseconds) and enhanced sampling simulations, we provide evidence that ligands selectively induce domain rearrangement, leading to interdomain contact. We use protein-protein interaction assays to provide experimental evidence of these interactions, identifying a critical role of the hinge in mediating interdomain contact. Our results illuminate previously unknown aspects of interdomain communication in FXR and provide a framework to enable characterization of other full length nuclear receptors.

## Introduction

Nuclear receptors are ligand-induced transcription factors that regulate the expression of target genes that are crucial in a myriad of biological processes, including development, metabolism reproduction, and cell cycle^1, 2^. In response to ligands, nuclear receptors bind specific DNA sequences and regulate gene programs. Members of this family share a common architecture, comprised of a disordered N-terminal domain, a DBD and an LBD, the latter two linked by a flexible hinge^3^. While the structure and activity of the two stably folded nuclear receptor domains (LBD and DBD) are well characterized, they are flanked by disordered regions whose functions are poorly understood. The difficulty of obtaining experimental structures of full-length nuclear receptor models has posed a significant limitation for structure-function analyses of disordered domains, as well as for deciphering the nature of interdomain crosstalk in receptors.

Of the 48 nuclear receptor genes in humans, full-length x-ray crystallographic structures have been reported for only five. This short list includes three RXR heterodimers^4–6^ and HNF4-^7^. A handful of low resolution cryo-electron microscopy structures have been published for RXR-VDR^8^, AR^9^, PR^10^, as well as EcR-USP^11^, the drosophila homolog of FXR-RXR^12^. An increasingly popular alternative for predicting full-length receptor structures is the use of integrated models that combine structural and biophysical methods with computational modeling. Thus far, structures of ER^13^and LRH-1^14^ have been predicted this way. Strikingly, very little structural overlap in quaternary architecture has been observed among existing models, suggesting the imprudence of generalizing across receptors or simply applying structural details of one receptor to another. More importantly, these observations highlight the dire need for new approaches that can facilitate the study of full-length structure in all nuclear receptors.

Our limited perspective on interdomain interactions in nuclear receptors also limits our understanding of long-distance (i.e. allosteric) communication between distant nuclear receptor domains. In well-studied mechanisms, ligand binding to the LBD induces conformational changes that influence the AF-2 surface to modulate coregulator recruitment^15^ . Ligands also induce promoter selective effects on transcription, as demonstrated in FXR^16^, suggesting that local information from the LBD may be transmitted to the DBD to influence interactions with DNA. While the molecular nature of this allosteric regulation is poorly understood, interdomain interfaces observed in existing full length nuclear receptors are proposed to act as conduits for communication. Indeed, evidence from biophysical experiments in multiple receptors suggests that ligands can both induce and modulate contact between LBD and DBD^17–19^.

Computational approaches hold immense promise for addressing both limitations described above, i.e., obtaining structural models of full-length receptors, and understanding the role of ligands in modulating their quaternary architecture. Because of powerful advances in homology modeling and machine-learning based structure prediction approaches^20,21^(e.g. Alphafold, RoseTTAa fold), it is now trivial to predict reliable structures of folded domains and generate starting configurations for disordered loops. Molecular dynamics (MD) simulations are a powerful tool for modeling dynamic behavior of disordered regions^22^, as well as for describing how small molecules modulate conformational dynamics in protein complexes^23^. The size of full-length receptors poses a challenge for detailed MD studies on timescales where interdomain motions can be observed. Recently, Chen et al. applied coarse-grained MD simulations to describe interdomain communication in human ER^13^, revealing that ligands uniquely modulate the hinge to facilitate interdomain communication. Here, we aim to present an atomistic perspective on how farnesoid X receptor (FXR) domains interact in the full structure and in the presence of diverse ligands.

In response to bile acid levels, FXR modulates the transcription of genes involved in lipid, bile acid, and glucose metabolism^24,25^. Because of its gene regulatory profile, FXR has received considerable attention as a drug target for several liver disorders and metabolic diseases^26–28^. Our primary goal is to describe DBD-LBD interdomain interactions in FXR. For simplicity, we have excluded the N-terminal from our full-length FXR (fl-FXR) model described here. We used homology modeling^19^ to predict the initial fl-FXR structure, based on liver X receptor beta (LXRβ, similarity: 54%) as a template. An ensemble of hinge conformations was generated using MD simulations, followed by clustering to identify optimal starting states. Enabled by the Anton2 Supercomputer^20^, we performed atomistic microsecond-scale MD simulations on multiple fl-FXR complexes, observing that ligands selectively induced rearrangement of FXR domains. To broadly sample fl-FXR dynamics, we employed accelerated MD simulations^21^, which permit prediction and visualization of interdomain interfaces in FXR. Finally, we use a protein-protein interaction assay to probe the predicted DBD-LBD interaction in FXR, revealing that the hinge plays an active role in mediating interdomain contact. These studies illustrate how MD simulations can generate accurate descriptions of full-length receptors, representing a crucial step towards the larger goal of characterizing interdomain allostery in the entire family of nuclear receptors.

### Prediction and optimization of structural models of full-length FXR

To overcome the limitations of obtaining experimental structures of nuclear receptors, we employed computational modeling to generate a structure of fl-FXR, i.e. FXR DBD and LBD (See Methods). LXRβ from PDB 4NQA^4^ was used as a template to obtain the initial domain arrangement of FXR DBD and LBD **(Fig. 1A)**. We used Modeller^29^ to predict three conformations of the flexible hinge **(Fig. 1B)**. To optimize the hinge conformation in preparation for longer simulations, we used accelerated MD simulations^30^ to explore the conformational space of the hinge. After obtaining 500 ns trajectories for each model, we combined and clustered the three trajectories to identify the top two conformations sampled by fl-FXR **(Fig. 1C**). We designate the two models as ‘extended’ and ‘compact’, based on the relative orientation of the DBD to the LBD **(Fig. 1D)**. In the extended model, the centers of mass of the domains are separated by ∼ 41.5 Å, with an angle of 104.8° between them. In the compact model, the domains are adjacent to one another with an angle of 62° between them. A comparison of existing full length nuclear receptor crystal structures shows that the relative DBD-LBD orientations in our extended and compact models lie within the range of interdomain angles observed in these experimental structures **(Fig. 1E)**. We used both fl-FXR models as the starting conformations for subsequent MD simulations.

**Figure 1.**
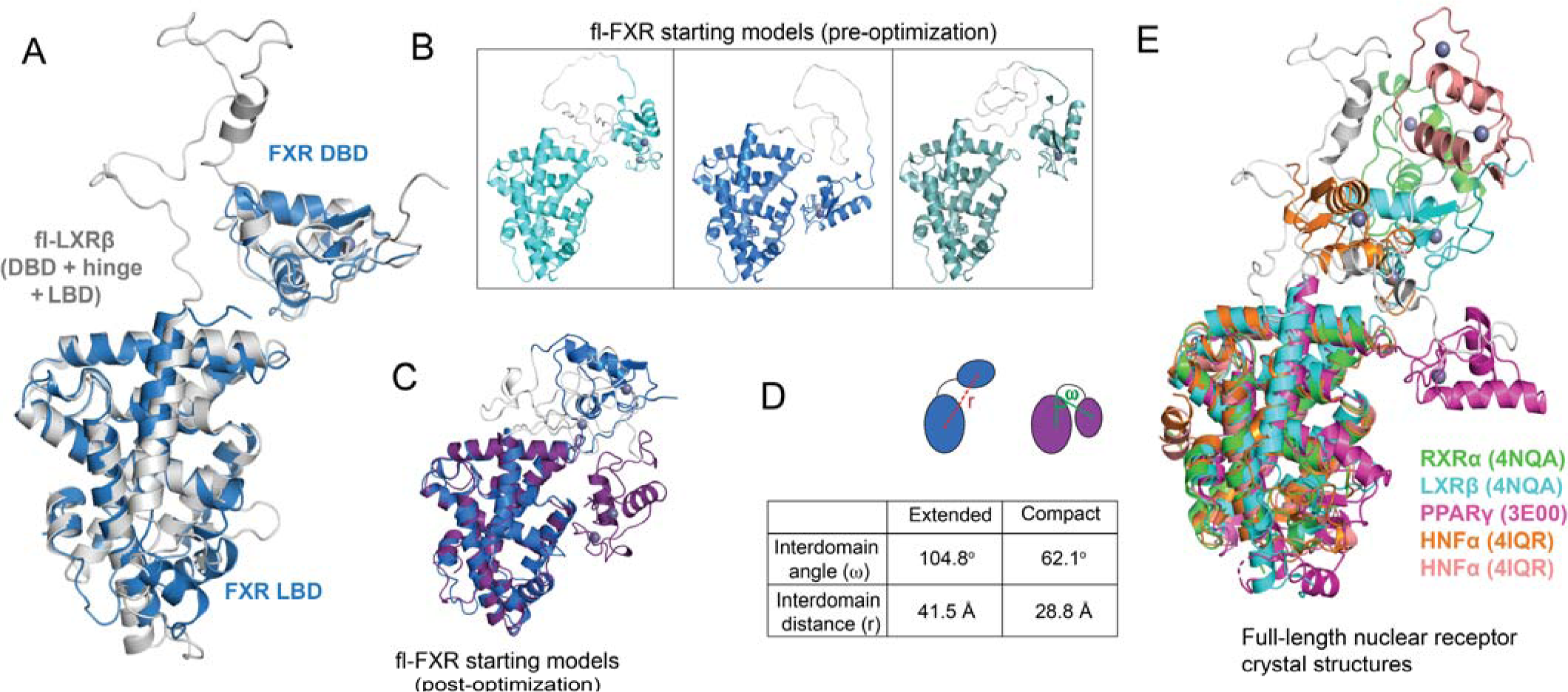
Structural models of full-length FXR. A) The fl-FXR model was generated by using full-length LXRβ (PDB 4NQA) as a template. Models of FXR DBD and LBD were aligned to LXRβ to predict initial arrangement of the domains. B) Modeller was used to insert the hinge region between the DBD and LBD. C) Two conformations of fl-FXR emerged following accelerated MD simulations to optimize the hinge conformation. D) Two fl-FXR conformations are designated as extended and compact. Interdomain angle (ω) and interdomain distance (r) parameters are illustrated on the models. E) Alignment of existing full length nuclear receptor crystal structures shows a range of interdomain (DBD-LBD) angles which encompass the angles of our extended and compact models.

### MD simulations capture domain rearrangement in full length FXR

To characterize our fl-FXR models in the presence of bound ligands, we constructed complexes with three FXR ligands: lithocholic acid (LCA), chenodeoxycholic acid (CDCA), and obeticholic acid (OCA). While LCA is a weak agonist/antagonist, both CDCA and OCA, the latter being a semisynthetic derivative of CDCA, are FXR agonists with calculated EC_50_ values of 10 µM and 99 nM, respectively^31^. Using both extended and compact fl-FXR models, we generated 10-20 microsecond long trajectories of all three ligand-bound forms, along with an apo (unliganded) state. To monitor conformational changes across the extended FXR trajectories, we quantified root mean square fluctuations (RMSF), interdomain distance (r) and interdomain angles (ω) for each complex. Among extended fl-FXR complexes, the most drastic change is observed in FXR-OCA, which shifts into a compact state (Fig. 2A), accompanied by a large decrease in both interdomain angle and distance (Fig. 2C). This mechanism is reminiscent of domain closure observed in enzymes such as adenylate kinase, which transitions from an open to closed state by forming interdomain salt-bridges and serves to bring substrates into close proximity for chemical reaction^32^. We note that the repositioning in FXR-OCA docks the DBD next to LBD helix 10 (H10), which is also the binding site of RXR in the FXR-RXR heterodimer **(Fig. S1)**, suggesting that this conformational state would preclude heterodimerization. FXR-CDCA also undergoes domain rearrangement but with smaller decreases in angle and distance. Apo-FXR and FXR-LCA both undergo minimal conformational changes (Fig. 2G, J), with small increases in interdomain angle but minor changes in DBD-LBD distance. Large fluctuations are observed in the flexible hinge, as well as in the DBD, consistent with the multiple unstructured loops in this domain (Fig. 2B, E, H, K).

**Figure 2:**
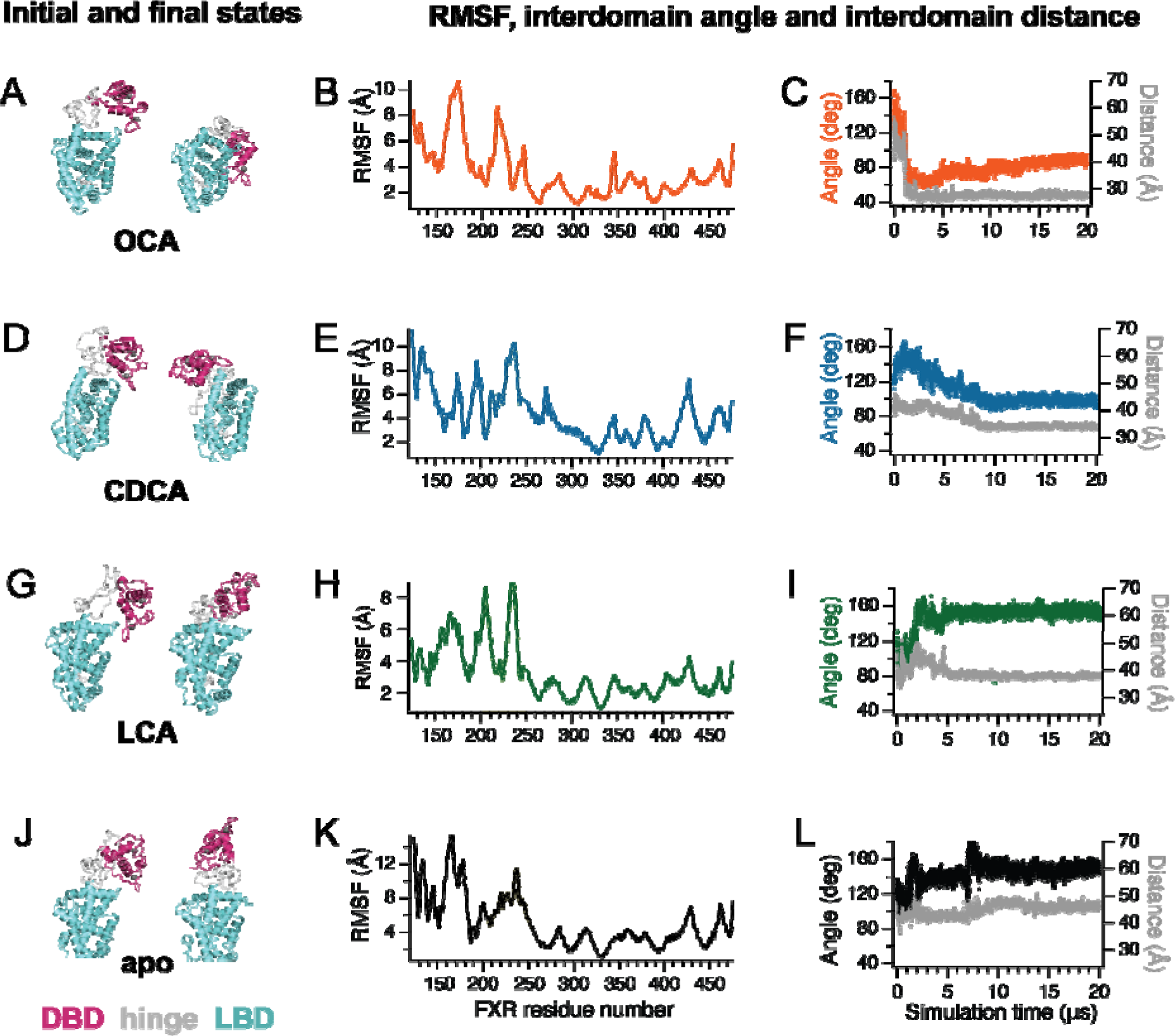
Analysis of extended fl-FXR MD simulations. Conformational changes, RMSF and changes in interdomain angle and distance are characterized for FXR-OCA (A-C), FXR-CDCA (D-F), FXR-LCA (G-I) and apo-FXR (J-L). DBD, hinge and LBD are colored magenta, grey and cyan respectively. FXR-OCA shifts from an extended to compact conformation over the 20 µs simulation (A, C). FXR-CDCA shifts into a partially compact conformation (D, F). FXR-LCA and apo-FXR do not undergo large conformational changes over the simulation (G, I, J, L). The highlighted region in RMSF plots is the hinge (B, E, H, K). For all complexes, largest fluctuations are observed in the DBD and hinge while the LBD remains stable.

We analyzed the compact fl-FXR complexes using the same methods **(Fig. S2)**. Similarly, the largest conformational change was observed in FXR-OCA, which switched from compact to an extended conformation. The other three complexes retained their compact state for the entirety of the simulation. As observed in extended FXR, large DBD and hinge fluctuations are observed while the LBD remains relatively stable.

These findings confirm the dynamic nature of fl-FXR and its ability to transition between both forms. This data also hints at the selectivity of ligand-mediated domain rearrangement, as we only observe it with the potent agonist (FXR-OCA) and partially in the weaker agonist complex (FXR-CDCA).

To determine whether binding affinity for FXR ligands are influenced by fl-FXR conformation, we used the MM-PBSA approach (see Methods) to calculate the affinity of both extended and compact fl-FXR for CDCA and ivermectin (IVM). Both ligands had higher affinity for the compact fl-FXR conformation than the extended structure **(Fig. S3)**. As these calculations measure enthalpy and do not account for entropic contributions, this result suggests that the proximity of the DBD/hinge and LBD in the compact state which stabilizes the LBD also enhances its interaction with ligands. We also note that binding energy is more favorable for agonist CDCA than for antagonist IVM. However, as binding affinity is not always correlated with activity^33^, we do not make any inference based on the observed trend.

### Prediction of interdomain interfaces

To broadly explore the conformational space of fl-FXR, we used accelerated MD simulations to achieve enhanced sampling of FXR complexed with CDCA, OCA, and synthetic ligands IVM and NDB. Using our extended FXR model, we obtained 8-10 trajectories (500 ns – 1.5 µs) of each complex, a total of 38 complexes. Interestingly, FXR adopts a range of compact and extended conformations, observed across all ligands regardless of their functional profile. This observation suggests that the bias potential that enhances sampling in accelerated MD may also obscure ligand-specific dynamics in fl-FXR. To identify the most prevalent conformational states adopted by ligand-bound FXR, we attempted to cluster the complexes using interdomain angle and distance **(Fig. S4)**. These two parameters alone were unable to distinguish between the relative spatial orientations of DBD and LBD. This is illustrated (Fig. 3A), using two FXR conformations with similar interdomain angles and distances, but with the DBD lying adjacent to different faces of the LBD. To account for these three-dimensional domain arrangements, we defined two new parameters: rotational angle (θ) and vertical define the plane Π. Rotational angle θ describes the angular displacement of the DBD the DBD relative to Π (see Methods). Importantly, θ identifies the interdomain interface, i.e. the LBD ‘face’ that interacts with the DBD. These faces include the H10/H7 face, H1-H3 face, H5/H7 edge, H12 edge and H9 edge (Fig. 3C). To characterize the range of conformations in fl-FXR, we plotted θ and d_v_ along with interdomain distance (r) (Fig. 3D), allowing us to identify three major clusters of DBD-LBD interfaces (DLI) in fl-FXR.

**Figure 3.**
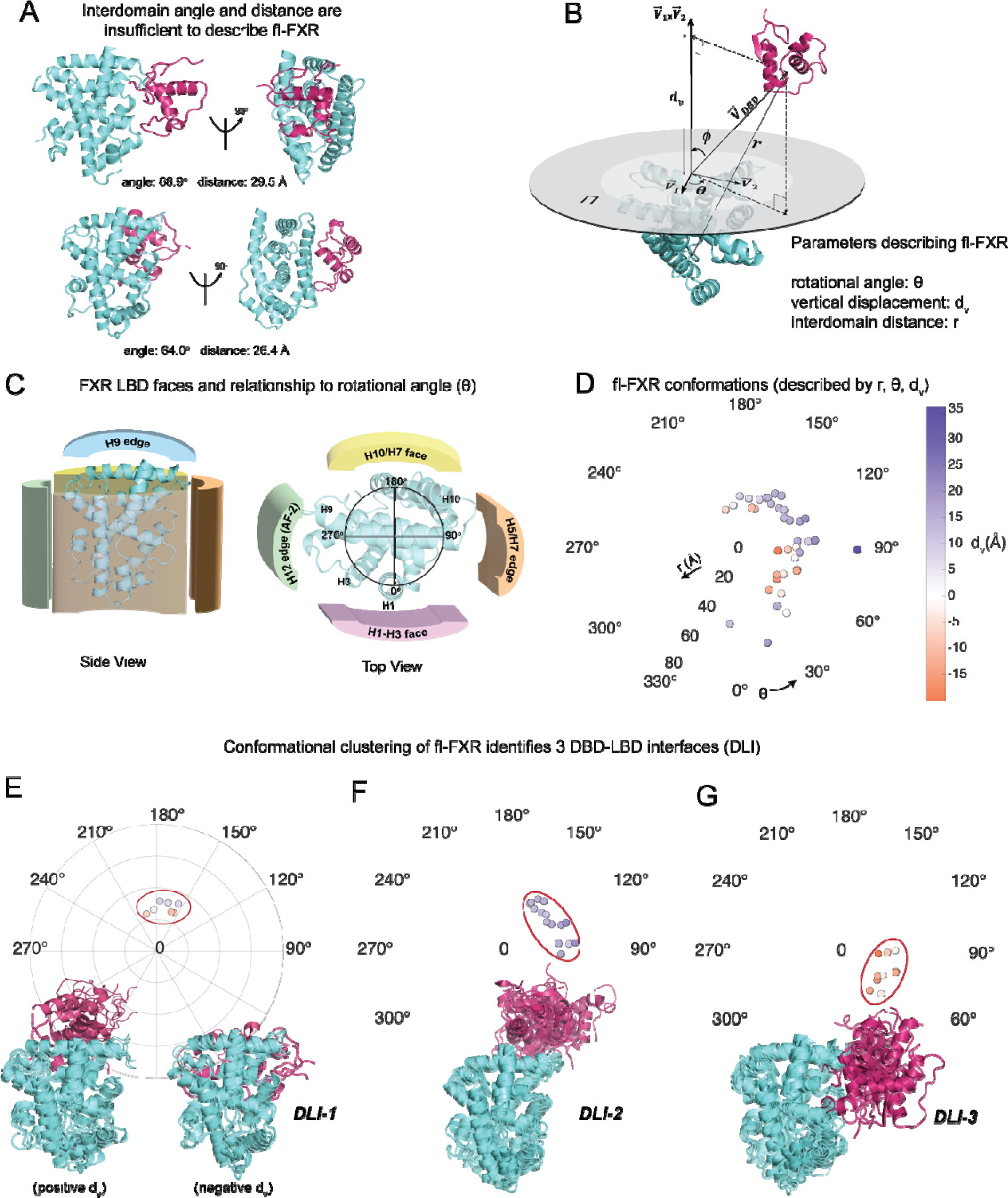
Conformational classification of fl-FXR. A) Two fl-FXR conformations with similar interdomain angle and interdomain distance have different 3D architectures. This illustrates that interdomain angle and distance are insufficient to describe the conformational ensemble of fl-FXR. B) We defined new parameters, namely rotational angle (θ) and vertical displacement (d_v_) to describe the 3D rotation of the DBD relative to the LBD. Combined with (r), these parameters can cluster the fl-FXR conformations from our simulations. C) To illustrate the rotational angle (θ), various interaction surfaces on the LBD are shown. For instance, a 180° rotation implies the DBD resides on the H10/H7 face, while a 90° rotation indicates that the DBD is located at the H5-H7 edge. D) Polar plot showing fl-FXR conformations described by rotational angle (θ), interdomain distance r (radially outward) and color-coded vertical displacement (d_v_). E-G) The conformations cluster into three groups (DLI-1, DLI-2, DLI-3), based on the interdomain interface (i.e. the LBD face/edge interacting with the DBD. E) In DLI-1, the interface is the H10/H7 LBD face. Two sub-groups are identified with positive or negative vertical displacement, respectively. F) The interdomain interface in DLI-2 is the H9 edge, all structures have positive d_v_. G) The interdomain interface in DLI-3 is the H5-H7 edge, all structures have negative d_v_.

In the first cluster, the DBD lies near the 180° rotation, positioned on the H10/H7 face of the LBD (Fig. 3E). Both positive and negative vertical displacements exist in the cluster, placing the DBD either above or below the plane Π. We designate this cluster as DBD-LBD interface 1 or DLI-1 (Fig. 3E). This positioning places the DBD in the RXR binding site, suggesting that this conformation can only exist in monomeric FXR. Of our 38 MD complexes, only 7 fall into the DLI-1 conformation, making it the smallest cluster. The second cluster, designated DLI-2, is defined by a rotational angle of 90-150° and a positive vertical displacement. This orientation places the DBD above the plane Π, at the H9 edge, similar to our previously designated ‘extended’ conformation (Fig. 1D).

The DLI-2 conformation accounts for 15 of 38 complexes (39%), making it the predominant fl-FXR conformation (Fig. 3F). The third cluster (DLI-3) accounts for 11 of 38 complexes (29%). DLI-3 is defined by a rotational angle of 30-90° and a negative vertical displacement, placing the DBD on the H5-H7 edge of the LBD (Fig. 3G). This conformation is similar to our ‘compact’ FXR state (Fig. 1D). While a few outliers exist outside of these 3 primary conformations, including two structures with rotational angle near 0, approaching the H1-H3 face, the DBD never approaches the H12 edge in our simulations. In summary, clustering of our MD simulations identifies 3 major conformations of fl-FXR: i) DLI-1 with the H10/H7 face as the interface between domains, ii) DLI-2 with the H9 edge as the interface, and iii) DLI-3 with the H5/H7 edge as the interface.

To compare structural predictions in fl-FXR with existing full length NR crystal structures, we calculated θ, d_v_ and r for RXRα-PPARγ (PDB 3E00), RXRα-LXRβ (PDB 4NQA), RXRα-RARβ (PDB 5UAN) and HNFα (PDB 4IQR)^4–6^ **(Fig. S5)**. Three of these occupy the DLI-2 conformation (HNF4α, LXRβ, RXRα) while two occupy DLI-3 (RARβ, PPARγ). The outliers (RXRα, HNF4α) have higher r values, conferring a hyper-extended conformation that is characteristic of structures that are part of a dimer and/or bound to DNA. The full-length LRH-1 model^34^ predicts a rotational angle of ∼ 0°, and would also be an outlier for the three DLIs in this work. Finally, to reveal whether ligands induce specific conformations in fl-FXR, we separated the complexes by ligand. While 6 of 10 OCA complexes occupy DLI-2, we observed more variation among DBD-LBD interfaces in the other ligand complexes **(Fig. S5)**. This observation confirms that while accelerated MD is useful for sampling the conformational space of fl-FXR, it is unable to resolve ligand-specific differences in interdomain conformation.

### Hinge-LBD salt bridges play a prominent role in stabilizing fl-FXR

Interdomain salt bridges are often important for stabilizing specific forms of multidomain proteins^35,36^. To characterize the critical salt bridges mediating interdomain interactions in fl-FXR as predicted by our simulations, we enumerated all salt bridges based on frequency of observation in the 38 fl-FXR trajectories. We focused on salt bridges present in at least 2 of the 38 complexes (summarized in **Table S1**). We grouped salt bridges as DBD-LBD (Fig. 4A), hinge-DBD (Fig. 4B) and hinge-LBD (Fig. 4C) salt bridges, identifying 17, 12 and 31 respectively that met our criteria. In addition to having the highest number of salt bridges among the three groups, the hinge-LBD group also contains the 8 most prevalent salt bridges (i.e. highest frequency among 38 trajectories).

**Figure 4.**
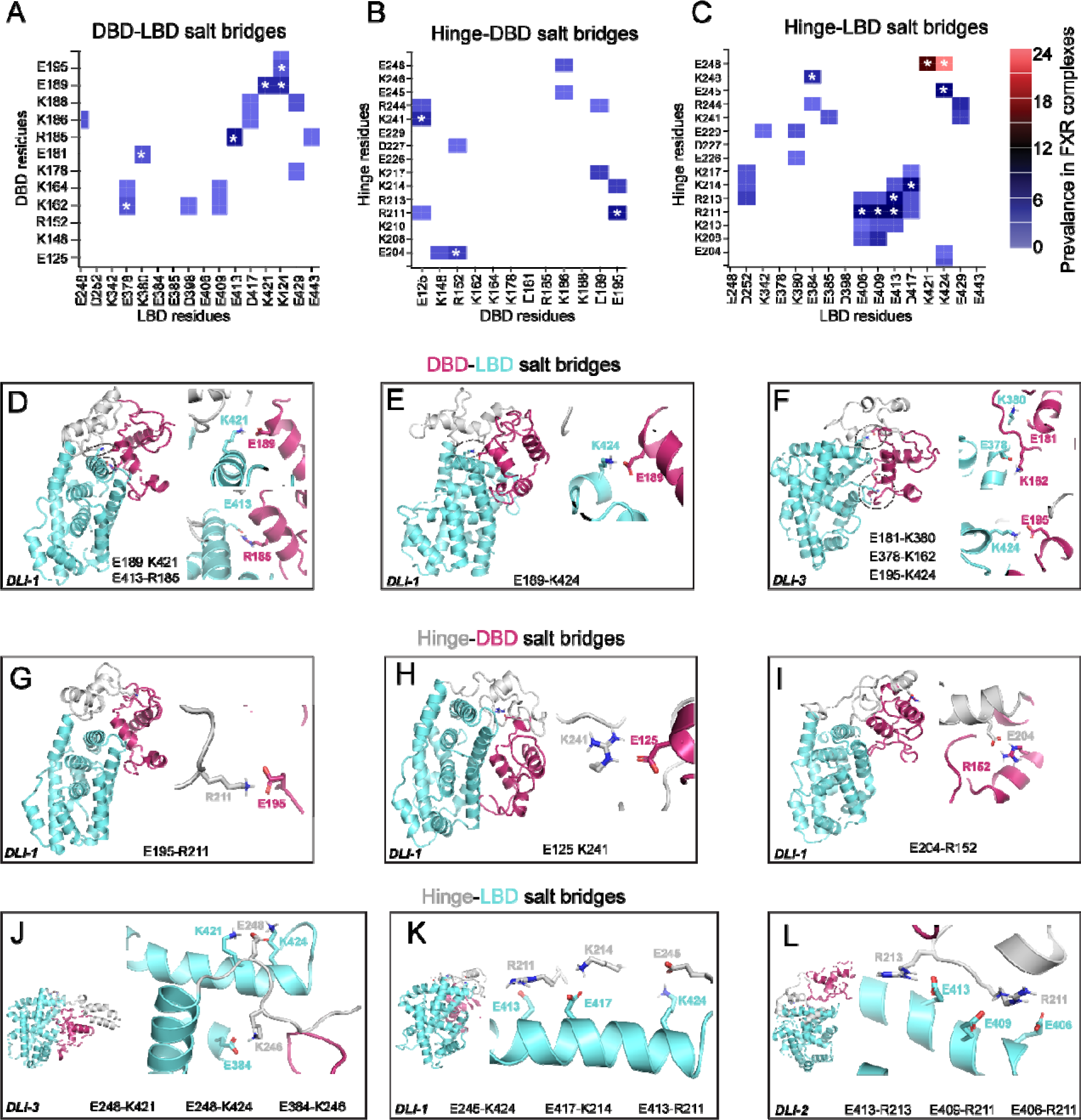
Interdomain salt bridges in fl-FXR. Salt bridges were identified from 38 accelerated MD trajectories of fl-FXR in various ligand bound states. Frequency of A) DBD-LBD, B) hinge-DBD, and C) hinge-LBD salt bridges are plotted as heat maps for comparison. Asterisks (*) indicate salt bridges illustrated in panels D-L. Of all three groups, hinge-LBD salt bridges are most prevalent across the 38 trajectories. D-E) DBD-LBD salt bridges are illustrated in DLI-1 and DL-3 conformations. G-I) Hinge-DBD salt bridges are illustrated in DLI-I conformations. J-L) Hinge-LBD salt bridges are illustrated in DLI-1, DLI-2 and DLI-3 conformations.

Interestingly, the three most prevalent DBD-LBD salt bridges occur between LBD H9 and DBD H2. The most frequent is between E413 and R185 and observed only in DLI-1 and DLI-2 complexes (Fig. 4D**, Table S1)**. Next is E189-K421, also observed in DLI-1 and DLI-2 conformations (Fig. 4D). The third most prevalent salt bridge involves K424, also on H9 but located at the C-terminal end, part of the H7-H5 edge (Fig. 4E). This position allows this salt bridge to stabilize DLI-3 conformations, as well as DLI-1. Less prevalent DBD-LBD salt bridges include E378-K162 and E181-K380, both involving LBD H7 residues and exclusively present in DLI-3 complexes (Fig. 4F). The E195-K424 salt bridge also stabilizes DLI-2 complexes via interaction with H9. In summary, DBD-LBD salt bridges are most likely to involve LBD H9 and occur in all three fl-FXR conformations, with larger representation in DLI-1 and DLI-2. Salt bridges involving H7 occur to a lesser extent and stabilize the DLI-3 state.

Unlike DBD-LBD salt bridges, hinge-DBD salt bridges are not associated with specific FXR conformations. The three most prevalent hinge-DBD salt bridges E195-R211, E125-K421 and E204-R152 (illustrated in Fig. 4 **G-I**) are observed across all 3 clusters **(Table S1)** which may indicate non-specificity in these interactions. Unlike with DBD-LBD interactions where the majority of DBD residues were from H2, the DBD residues implicated in the top three DBD-hinge salt bridges are from different structural motifs of the DBD, suggesting that there is no region particularly favored for these interactions. In summary, DBD-LBD salt bridges are likely to be non-specific and not making major contributions to fl-FXR architectures.

The hinge-LBD salt bridges are significantly more prevalent than both DBD-LBD and hinge-DBD salt bridges (Fig. 4C). Further emphasizing the importance of LBD H9, we observe that the 8 most prevalent of all salt-bridges are between the hinge and H9, encompassing conformations across the three clusters (Fig. 4 **J-L, Table S1).** The two most prevalent salt bridges are E248-K424 and E248-K421(Fig. 4J), present in 24 and 16 of our 38 trajectories. This analysis suggests that hinge-LBD interactions may be of particular importance for fl-FXR, more so than LBD-DBD or hinge-DBD interactions. Our simulations also strongly implicate LBD H9 in critical interdomain interactions.

### Experimental validation of ligand-induced interdomain contact in FXR

To test the hypothesis that FXR DBD and LBD form direct interdomain contacts, we employed a mammalian two hybrid cellular assay. This commonly used assay for protein-protein interaction has been used to demonstrate interactions between the N-terminal domain and LBD of steroid receptors^37–39^. We prepared hybrid protein constructs by fusing the FXR LBD to the Gal4-DBD (*Gal4DBD-(FXR-LBD)*) and the FXR DBD to the VP16 activation domain (*VP16-(FXR-DBD)*) (Fig. 5A). To probe the role of the hinge in DBD-LBD interactions, we designed additional hybrid constructs attaching the hinge separately to the LBD (*Gal4DBD-(FXR-hinge-LBD)*) and DBD (*VP16-(FXR-DBD-hinge)*) (Fig. 5A). To quantify background activity, we used VP16 only without the fused FXR DBD (*VP16-control*). We observed background luciferase activity resulting from the inherent transcriptional capability of the Gal4DBD-(FXR-LBD) construct.

**Figure 5.**
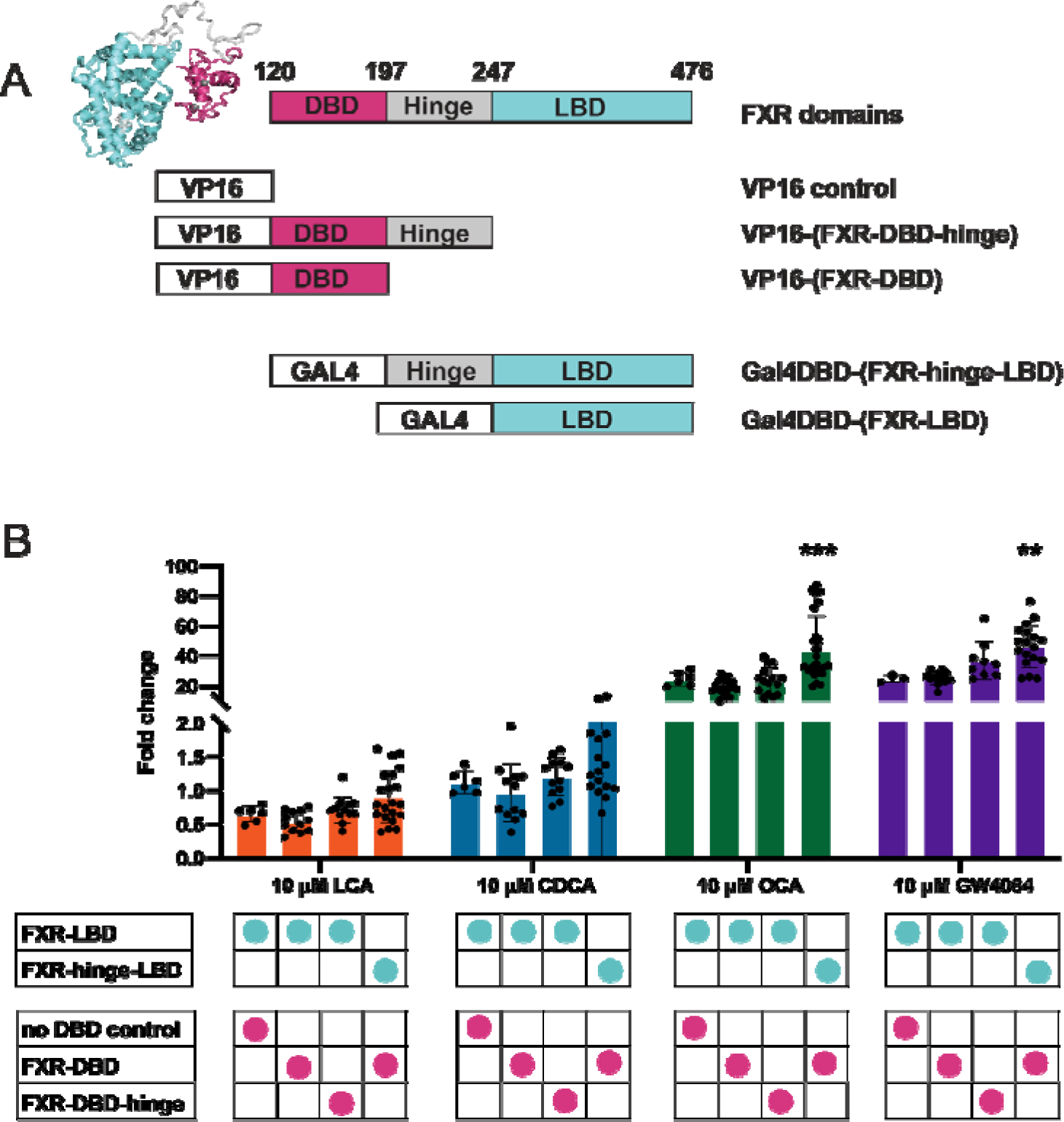
Experimental validation of interdomain interaction in FXR. A) Four mammalian two hybrid fusion constructs were prepared for this study. The VP16 activation domain was fused to the FXR DBD (residues 120-196) and DBD plus hinge (residues 120-244). The Gal4DBD was fused to the FXR LBD (residues 247-476) and to the hinge plus LBD (residues 197-476). An unfused VP16 protein was used for experimental controls. B) Transcriptional activity of the luciferase gene under control of the UAS promoter is used to measure the interaction between Gal4DBD and VP16 fusion constructs. Data are reported as fold changes over control with no ligand (DMSO only). For all four ligands, transcription is measured under four conditions: LBD + no DBD control, LBD + DBD, LBD + DBD-hinge, and hinge-LBD + DBD. Background activation is observed in the control (no DBD) condition. No significant increase above background is observed for any ligand in the DBD + LBD condition. Similarly, no significant increase above background is observed in the DBD-hinge + LBD condition. A significant increase is only observed in the DBD + hinge-LBD condition, indicative of an interaction between the two fusion proteins.

When VP16-(FXR-DBD) was introduced, no increase in signal was observed, indicating the absence of a direct interaction between FXR-LBD and DBD. Next, we tested our VP16-(FXR-DBD-hinge) construct, to determine whether the addition of the hinge introduces interdomain contact. We observed a small but significant increase only with potent nonsteroidal FXR agonist GW4064, but not with other ligands (Fig. 5B). Finally, we tested the interaction between the Gal4-(FXR-hinge-LBD) construct and VP16-(FXR-DBD), observing an increase in luciferase activity in both OCA and GW4064 but not in weaker ligands LCA and CDCA (Fig. 5B). To provide further support for this ligand-specificity, we increased the concentration of LCA and CDCA to 100 µM and still did not observe a significant difference between the LBD-only and hinge-LBD constructs **(Fig. S5)**. To confirm that the increased luciferase signal results from hinge-induced contact between LBD and DBD, we tested the interaction between the hinge-LBD construct and the VP16-control with no DBD present (Fig. 5B). We also observed that the presence of the DBD significantly increases transcription, confirming the interdomain interaction **(Fig. S6)**. These results suggest an important role for the hinge in mediating contact between LBD and DBD, as well as the ligand-specific nature of this interaction.

## DISCUSSION

Understanding the dynamic nature of full-length nuclear receptors at a molecular level has been an elusive goal. Existing crystal structures largely present full-length nuclear receptors in an extended conformation, often DNA-bound or dimerized. Nonetheless, biophysical experiments provide evidence of interdomain DBD-LBD contact, suggesting that nuclear receptors are dynamic and capable of shifting from extended to compact forms. Our knowledge of full-length nuclear receptor structure is limited, and to a greater extent, so is our understanding of full-length receptor dynamics. Here, we use computational modeling to predict initial conformations of fl-FXR for subsequent simulation studies. Our simulations indicate that FXR domains rearrange between extended and compact states, similar to domain closure observed in enzymology which is critical for positioning substrates for catalysis. Further, we show via simulations that domain rearrangement is ligand-modulated in fl-FXR, corroborating previous studies where interdomain contacts have been observed in full-length nuclear receptors ^13,18,40,41^. Mammalian two-hybrid experiments confirm that the FXR LBD and DBD interact only when the hinge is included. The DBD-hinge construct did not interact with the LBD, suggesting that the presence of the hinge alone is not sufficient. It is only when attached to the LBD that the hinge mediates interdomain contact. Thus, the hinge is not just a linker, but plays an active role in mediating interdomain contact. Notably, the hinge-LBD salt bridges are the most prevalent interdomain salt bridges observed in our MD simulations. This result is consistent with previous studies where an interdomain linker region is demonstrated to mediate allosteric communication between folded domains^42–44^.

Consistent with predictions from simulations, we also observe that this interdomain contact is ligand-modulated, as it only observed in potent agonists OCA and GW4064 and not in weaker ligands, even when 100 µM CDCA or LCA is added. Based on these observations, we propose a mechanism whereby ligand binding propagates a conformational change to the hinge, enabling interaction with the DBD, possibly by creating the binding site. We posit that weaker ligands are unable to induce this conformational change in the hinge, either to the same degree or at all. Subsequent studies will aim to characterize the thermodynamics of the interdomain contact in FXR. While the interaction between LBD, hinge and DBD in the compact state is stabilized by salt bridges and other noncovalent forces, there is likely a tug-of-war between enthalpy and entropy, as the ‘closed’ state is less entropically favorable than extended fl-FXR. It is possible that the potency of a ligand is related to the ΔΔG of binding compact versus extended FXR.

While our studies provide insight on the physical nature of interdomain communication in fl-FXR, they do not inform about the physiological relevance or timing of domain rearrangement in fl-FXR signaling. This type of rearrangement seems most plausible in a monomeric receptor, which is a transcriptionally active form for FXR^45^. Future studies will investigate the dynamical nature of the FXR-RXR heterodimer to assess whether domain rearrangement occurs in the dimeric state. Domain rearrangement may also occur prior to FXR-RXR dimerization, a scenario which suggests both a steep increase in entropic penalty and a more favorable enthalpy due to inter-receptor interactions. To the best of our knowledge, this study reports the first observation of an open (extended) to closed (compact) domain rearrangement in a nuclear receptor. Similar domain closures have only been reported in enzymes. As nuclear receptors share a conserved structure and mechanism, we anticipate that similar ligand-modulated domain rearrangement will be observed in other receptors.

## Materials and Methods

### Protein sequence and structures

To construct a model of full-length FXR (fl-FXR), we first used Modeller V9.23 to build a model of FXR DBD (residues D124-Q476, Uniprot Q96RI1-1), which had not been crystallized at the time. This structure has since been solved^46^and aligns with our model to RMSD < 2 Å. As a template, we used retinoic acid receptor alpha DBD taken from chain A of PDB 1DSZ (RARα:FXR DBD sequence similarity = 68%). Together with the FXR LBD from PDB 6HL1, a homology model was created by aligning the two domains to the corresponding domains of full-length LXRβ (LXRβ-FXR similarity: 54%) obtained from PDB 4NQA. Modeller was then used to predict three conformations of the interdomain hinge as a starting point for further optimization via simulations.

### Classical MD simulations

Extended MD simulations were performed on Anton 2^47^. For binding energy calculations, triplicate 500 ns simulations were obtained using Amber18^48^ with GPU acceleration^49^. Antechamber^50^ from AmberTools^51^ was used to parameterize FXR ligands. The ff14SB forcefield^52^ and Generalized Amber Forcefield2^53^ were used for proteins and ligands, respectively. Complexes prepared for simulation on Anton 2 were solvated in a cubic box (103 x 98 x77 Å^3^) of TIP3P water^54^, with sodium and chloride ions added to reach a concentration of 150 mM NaCl. Complexes prepared for energy calculations were solvated in an octahedral box with a 10 Å buffer.

All complexes were minimized, heated and equilibrated using the Amber18. Minimization was performed in four steps: i) with 500 kcal/mol. Å^2^ restraints on solute atoms, ii) 100 kcal/mol.Å^2^ restraints on solute atoms, iii) 100 kcal/mol. Å^2^ restraints on ligand atoms only, and iv) with no restraints on any atoms. Each minimization step utilized 5000 steps of steepest descent followed by 5000 steps of conjugate gradient. Heating to 300 K was performed using a 100-ps NVT simulation with 5 kcal/mol.Å^2^ restraints on all atoms. Pre-equilibration was performed in three 10-ns steps: i) with 10 kcal/mol.Å^2^ restraints on solute atoms, ii) with 1 kcal/mol.Å^2^ restraints on solute atoms, iii) with 1 kcal/mol.Å^2^ restraints on ligand atoms. After restraints were removed, Anton complexes were equilibrated for 50 ns before transferring to Anton 2 for extended MD. Complexes for energy calculations were simulated for 500 ns in triplicate. For all simulations, a 2-fs timestep was used with SHAKE. To evaluate the long-range electrostatics with particle mesh Ewald^49^ and Van-Der Waals forces, a 10-Å cutoff was used. CPPTRAJ^55^, MDtraj^56^ and MDAnalysis software were used to analyze RMSF and Salt-bridges. Binding free energy calculations were performed using the MM-PBSA^57^ method in the AMBER.

### Accelerated MD simulations

To achieve enhanced conformational sampling of these multidomain proteins, we employed accelerated MD simulations (aMD)^30,58^, a type of enhanced sampling method where the potential energy surface is modified by applying a boost potential when the potential lies below a certain minimum. Thus, it allows the simulation to sample different parts of the energy surface faster. We have applied a dual boost potential in our study. The boost potential (ΔV) is calculated as follows:

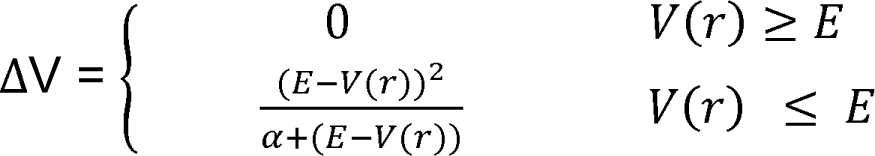

The method discussed in the published protocol^59^ was followed here to calculate the different parameters needed for aMD simulations. All aMD simulations were done using AMBER20 software. VMD^60^ and PyMol^61^ were used to visualize the protein structures and simulation trajectories. ProteinTools was used to visualize salt-bridges and hydrogen-bonds.

### Cluster Analysis

We used CPPTRAJ to calculate the vectors 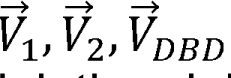, angle Φ and interdomain distance *r* for each complex. For fl-FXR model, the alpha carbon (Cα) of D417 was chosen as the origin of all vectors. V1 terminates at Cα-P251, V2 terminates at Cα-K424 and 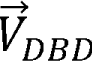 terminates at the center-of-mass of the DBD (residues 124-196). Angle Φ is the angle that 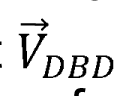 makes with 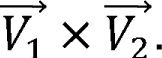. Interdomain distance is the distance between the center-of-mass of the DBD (residues 124-196) and center-of-mass of the LBD (residues 251-476). The plane ∠ was defined as the plane spanned by 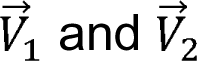. Vector algebra was then used to calculate the parameters θ and *d_v_* according to the following steps:

1. Gram-Schmidt orthonormalization of 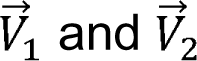 to construct orthonormal basis vectors 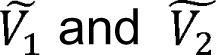 that span the plane ∠.

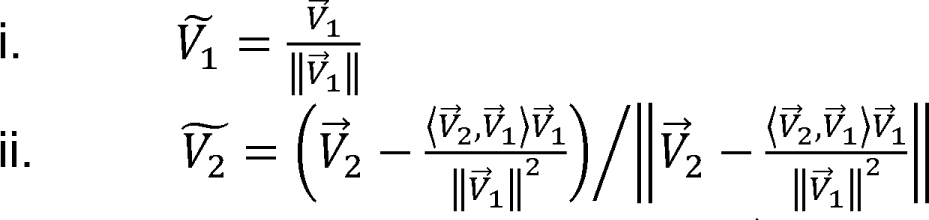

where 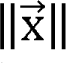 represents the norm of 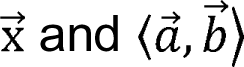 represents the inner product of 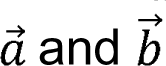.

1. Defined i. 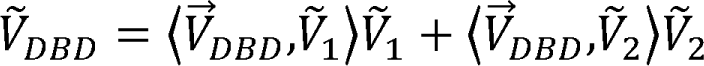

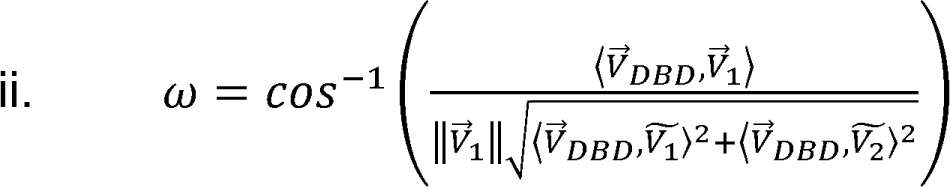

Then, θ was defined as,

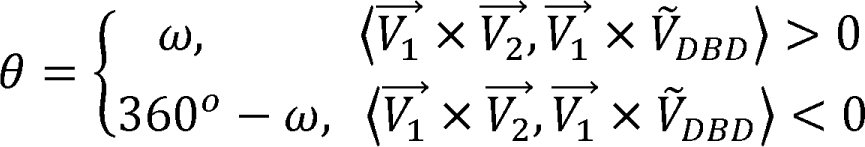

where 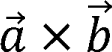 represents the vector product of 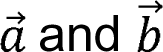.

(The sense of rotation of 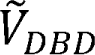 is counter-clockwise with respect to 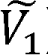)

3. 

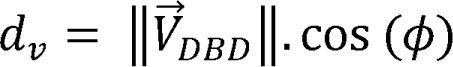

The above parameters were described in a similar fashion for the other full-length crystal structures **(Table S2)**.

### Mammalian two-hybrid assay

Mammalian two-hybrid assays were performed following the instruction of the CheckMate^TM^ Mammalian Two-Hybrid System (Promega). Constructs were synthesized by Genscript (Fig. 5A). The FXR LBD (residues 247-476) and the hinge region (residues 197-244) were cloned into the mammalian Gal4DBD fusion vector pBIND (Promega), to give Gal4DBD-(FXR-LBD) and Gal4DBD-(FXR-hinge-LBD), respectively. The FXR DBD (residues 120-196) and the hinge region (residues 197-244) were cloned into the pACT plasmid (Promega) to generate fusion proteins with the VP16 activation domain, resulting in VP16-(FXR-DBD) and VP16-(FXR-DBD-hinge). The amino acid numbering is in consistent with the sequence found in Uniprot Q96RI1-1.

HeLa cells were cultured in Minimum Essential Medium alpha (MEM) supplemented with 10% fetal bovine serum (FBS) and 1% of L-glutamine. Cells were plated 10,000 cells/well in 96-well, clear flat, bottom cell culture plate. Co-transfection was performed with equal amount of 5ng of each pBIND and pACT constructs, along with the 50ng reporter plasmid pG5luc, which contains the UAS response element and encodes firefly luciferases, using Fugene HD (Promega). Controls included wells with empty pACT vector, VP16 control. Transfection was repeated at least three times. After 24 hr incubation at 37°C in a 5% CO_2_ incubator, DMSO or test ligands were added at a final concentration of 1.3%, 10µM or 100µM, as appropriate.

After another 24hr of incubation, Firefly luciferase activity and Renilla luciferase activity were measured using the Dual-Glo kit (Promega) using a SpectraMax iD5 plate reader. Fold activation was represented as normalized luciferase over DMSO-treated control. A two-way ANOVA was used to evaluate the variance among the groups, followed by Tukey’s multiple comparison test to access the differences between specific pairs of means. All statistical analyses were performed using GraphPad Prism V10 software.

## Acknowledgments

Anton 2 computer time was provided by the Pittsburgh Supercomputing Center (PSC) through Grant R01GM116961 from the National Institutes of Health. The Anton 2 machine at PSC was generously made available by D.E. Shaw Research. C.D.O. is funded by an NSF award (CAREER: 2144679).

## Notes

### Competing Interest Statement

The authors have declared no competing interest.

### Summary of Updates

Additional analysis have been incorporated. A new figure (Figure 3) has been added to describe full length FXR conformations. Figure 4 has been updated to describe specific salt bridges stabilizing interdomain interactions in FXR. Supplemental information has been updated.

