## Supporting Information for "Nuclear receptor interdomain communication is mediated by the hinge with ligand specificity"

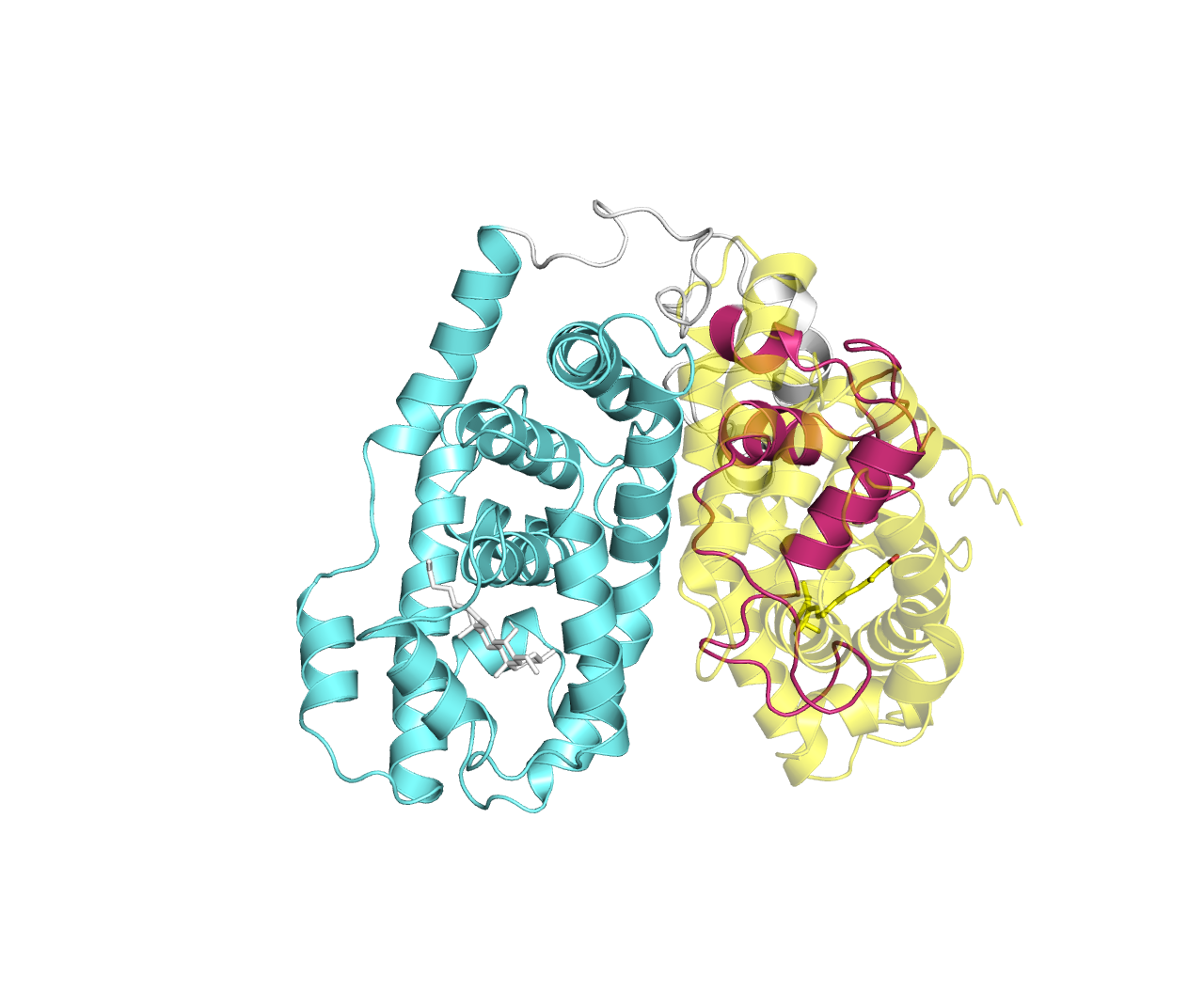

**Figure S1.** FXR-OCA transitions to a compact state which places the FXR DBD (magenta) in the binding site for RXR**α**, shown in yellow for reference. FXR LBD is shown in cyan.

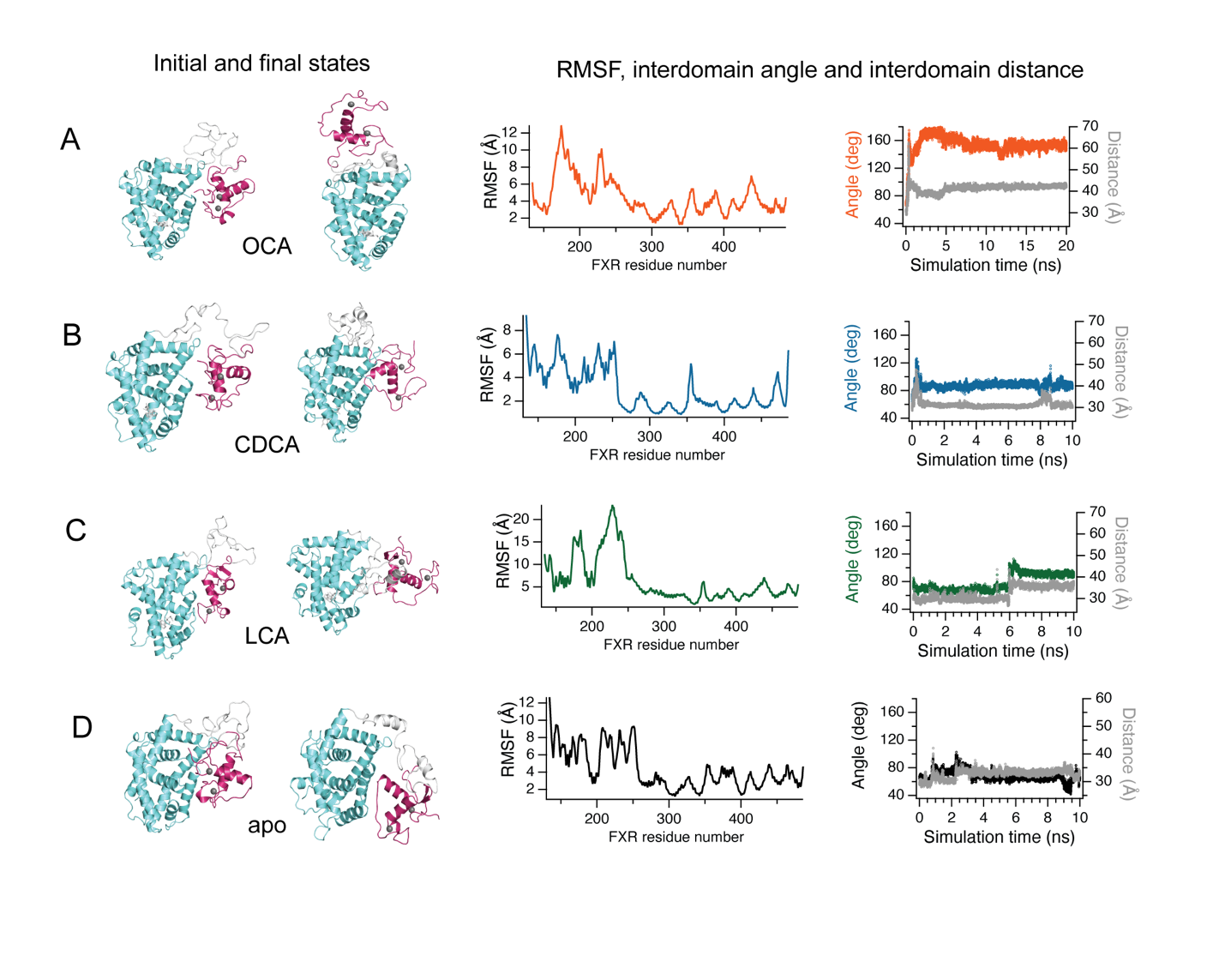

**Figure S2.** Analysis of compact fl-FXR MD simulations. Conformational changes, RMSF and changes in interdomain angle and distance are characterized for (A) FXR-OCA (B) FXR-CDCA (C) FXR-LCA and (D) apo-FXR. DBD, hinge and LBD are colored magenta, grey and cyan respectively.

| **Ligand** | **Binding free energy (Kcal/mol)** |
| --- | --- |
| FXR-CDCA (extended) | -21.87 ±1.009 |
| FXR-CDCA (compact) | -26.32 ± 1.149 |
| FXR-IVM (extended) | -2.74  ±  0.850 |
| FXR-IVM (compact) | -11.29 ± 0.805 |

**Figure S3.** Ligand binding energy of FXR complexes in extended and compact states. For both ligands, binding is more favorable to the compact state.

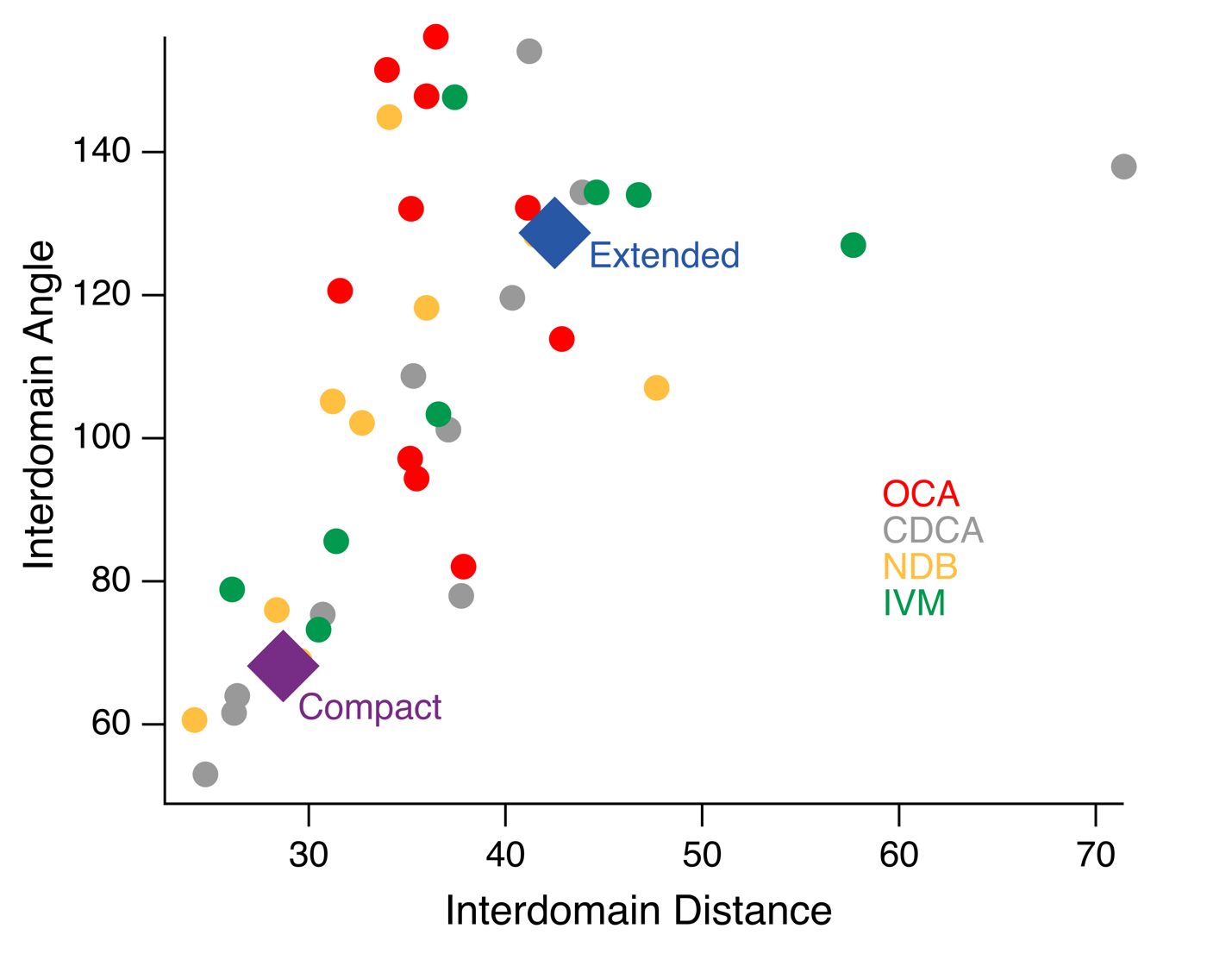

**Figure S4**. Analysis of fl-FXR conformations using interdomain angle and distance. These two parameters alone are limited for clustering, as they do not reveal any patterns among FXR conformations.

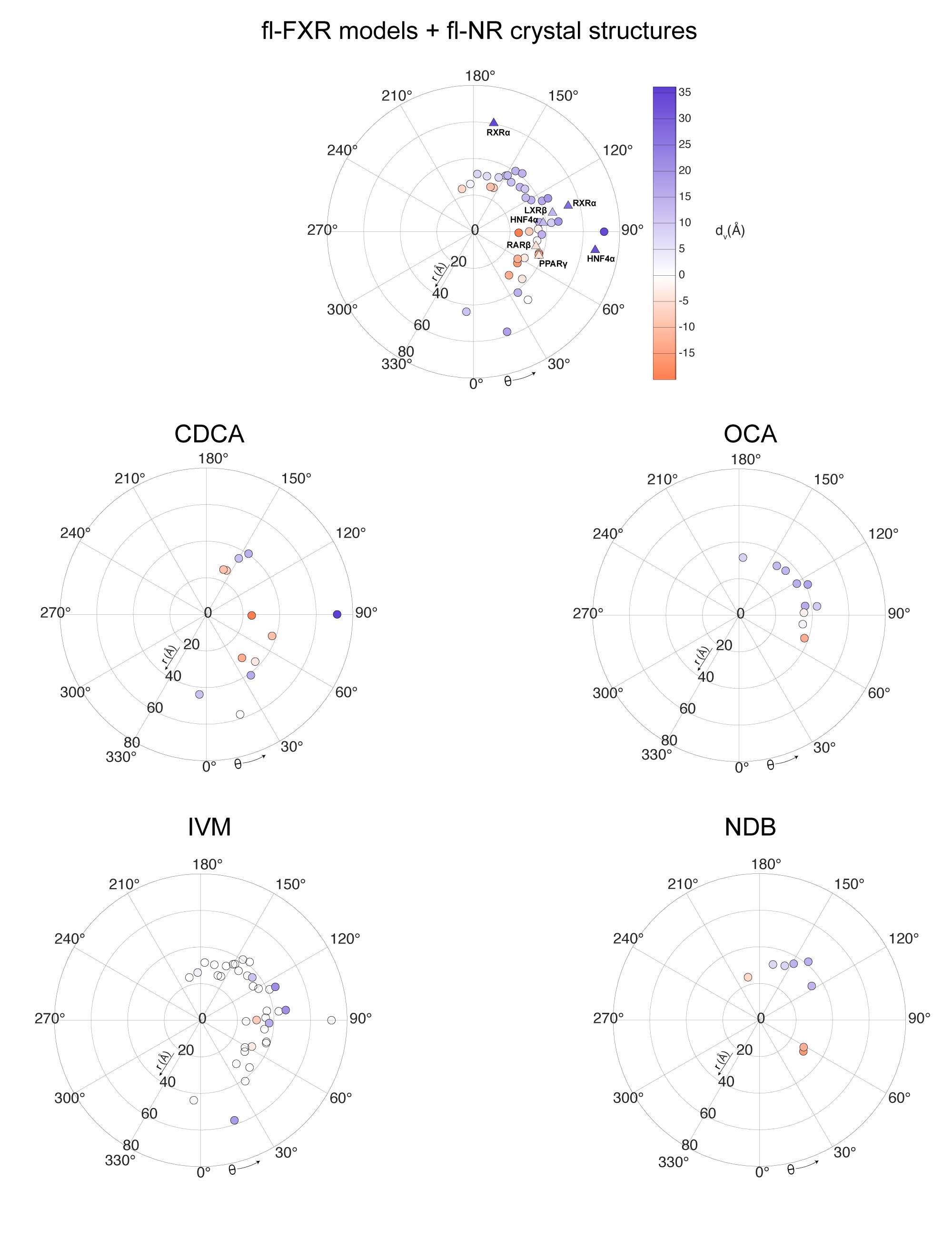

**Figure S5.** fl-FXR and fl-NR conformations described by rotational angle (θ), vertical displacement (d_v_) and interdomain distance (r).

**Table S1.** Interdomain salt bridges stabilizing fl-FXR conformations

| **Acceptor** | **Donor** | **DLI-1** | **DLI-2** | **DLI-3** | **Outlier** | **Total** |
| --- | --- | --- | --- | --- | --- | --- |
| **Hinge-LBD** |  |  |  |  |  |  |
| E248 | K424 | 4 | 9 | 9 | 2 | 24 |
| E248 | K421 | 3 | 8 | 4 | 1 | 16 |
| E409 | R211 | 1 | 7 |  | 0 | 8 |
| E413 | R211 | 1 | 6 | 1 | 0 | 8 |
| E245 | K424 | 3 | 2 |  | 1 | 7 |
| E413 | R213 | 1 | 5 |  | 1 | 7 |
| D417 | K214 | 2 | 3 | 1 | 1 | 7 |
| E406 | R211 | 1 | 6 |  | 0 | 7 |
| E384 | K246 |  | 1 | 4 |  | 5 |
| E429 | R244 | 1 | 3 |  | 1 | 5 |
| E409 | K208 | 3 | 2 |  |  | 5 |
| E429 | K241 | 3 | 1 |  |  | 4 |
| D252 | R213 |  | 2 | 1 | 1 | 4 |
| E406 | K208 | 1 | 1 |  | 1 | 3 |
| D417 | K213 |  | 3 |  | 0 | 3 |
| D252 | K214 |  | 2 |  | 1 | 3 |
| E229 | K380 |  | 1 | 1 | 1 | 3 |
| E413 | K214 | 1 | 1 |  | 1 | 3 |
| E409 | K213 |  | 2 |  | 1 | 3 |
| E413 | K210 | 1 | 1 |  | 1 | 3 |
| D252 | K217 |  | 2 |  | 1 | 3 |
| E406 | K213 |  | 1 | 1 | 1 | 3 |
| D417 | R211 | 2 |  | 1 |  | 3 |
| E385 | K241 |  |  | 3 |  | 3 |
| E409 | K210 |  | 2 |  | 1 | 3 |
| E204 | K424 |  | 1 | 1 |  | 2 |
| E384 | R244 |  | 1 | 1 |  | 2 |
| E406 | K210 |  | 1 |  | 1 | 2 |
| E226 | K380 |  | 1 | 1 |  | 2 |
| E229 | K342 |  |  | 1 | 1 | 2 |
| D417 | K217 | 1 |  |  | 1 | 2 |
| **Hinge-DBD** |  |  |  |  |  |  |
| E195 | R211 | 3 | 2 | 1 |  | 6 |
| E125 | K241 |  | 5 |  |  | 5 |
| E204 | R152 | 1 | 3 |  |  | 4 |
| E195 | K214 |  | 2 | 1 | 1 | 4 |
| E189 | K217 | 1 | 2 | 1 |  | 4 |
| E204 | K148 | 2 |  | 1 |  | 3 |
| E125 | R244 |  | 2 |  | 1 | 3 |
| E125 | R211 |  | 2 |  |  | 2 |
| E245 | K186 |  | 1 | 1 |  | 2 |
| E189 | R244 |  | 1 | 1 |  | 2 |
| D227 | R152 | 1 | 1 |  |  | 2 |
| **DBD-LBD** |  |  |  |  |  |  |
| E413 | R185 | 3 | 3 |  |  | 6 |
| E189 | K421 | 1 | 3 |  | 1 | 5 |
| E189 | K424 | 2 | 1 | 2 |  | 5 |
| E181 | K380 |  |  | 3 |  | 3 |
| E378 | K162 |  |  | 3 |  | 3 |
| E195 | K424 |  | 2 | 1 | 0 | 3 |
| E429 | K188 |  | 2 | 1 | 0 | 3 |
| E248 | K186 |  | 1 | 1 | 1 | 3 |
| D417 | K186 |  | 2 |  |  | 2 |
| D417 | K188 | 2 |  |  |  | 2 |
| E378 | K164 |  |  | 2 |  | 2 |
| E429 | K178 |  | 1 |  | 1 | 2 |
| E443 | R185 | 2 |  |  |  | 2 |
| D398 | K162 | 2 |  |  |  | 2 |
| E409 | K162 | 2 |  |  |  | 2 |
| E409 | K164 | 1 | 1 |  |  | 2 |

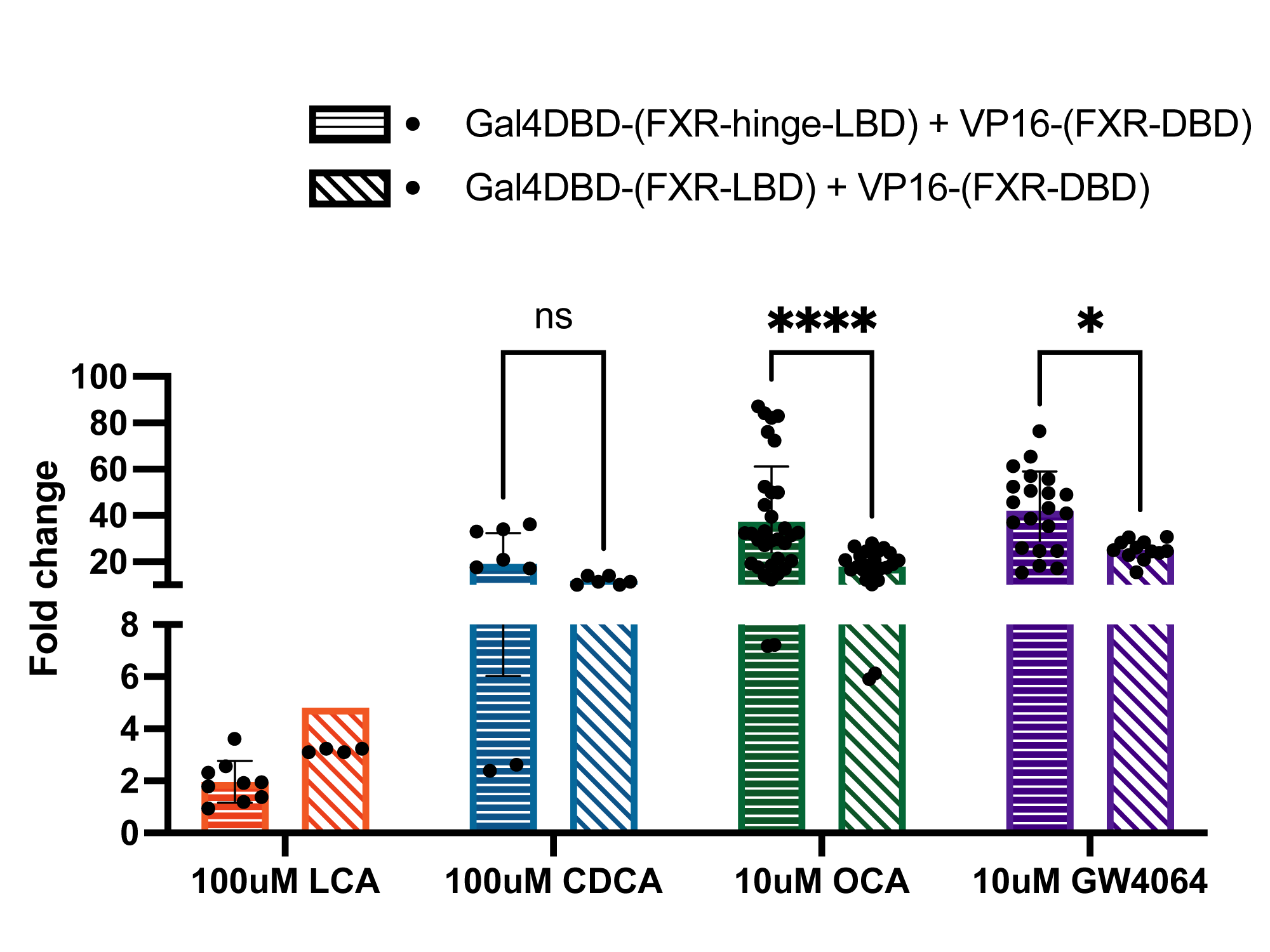

**Figure S6**. Mammalian two hybrid assays to compare protein-protein interaction between FXR-hinge-LBD and DBD vs LBD only and DBD. Even at 100 µM concentrations of the weak ligands CDCA and LCA, no significant increase in activity is observed in the presence of the hinge.

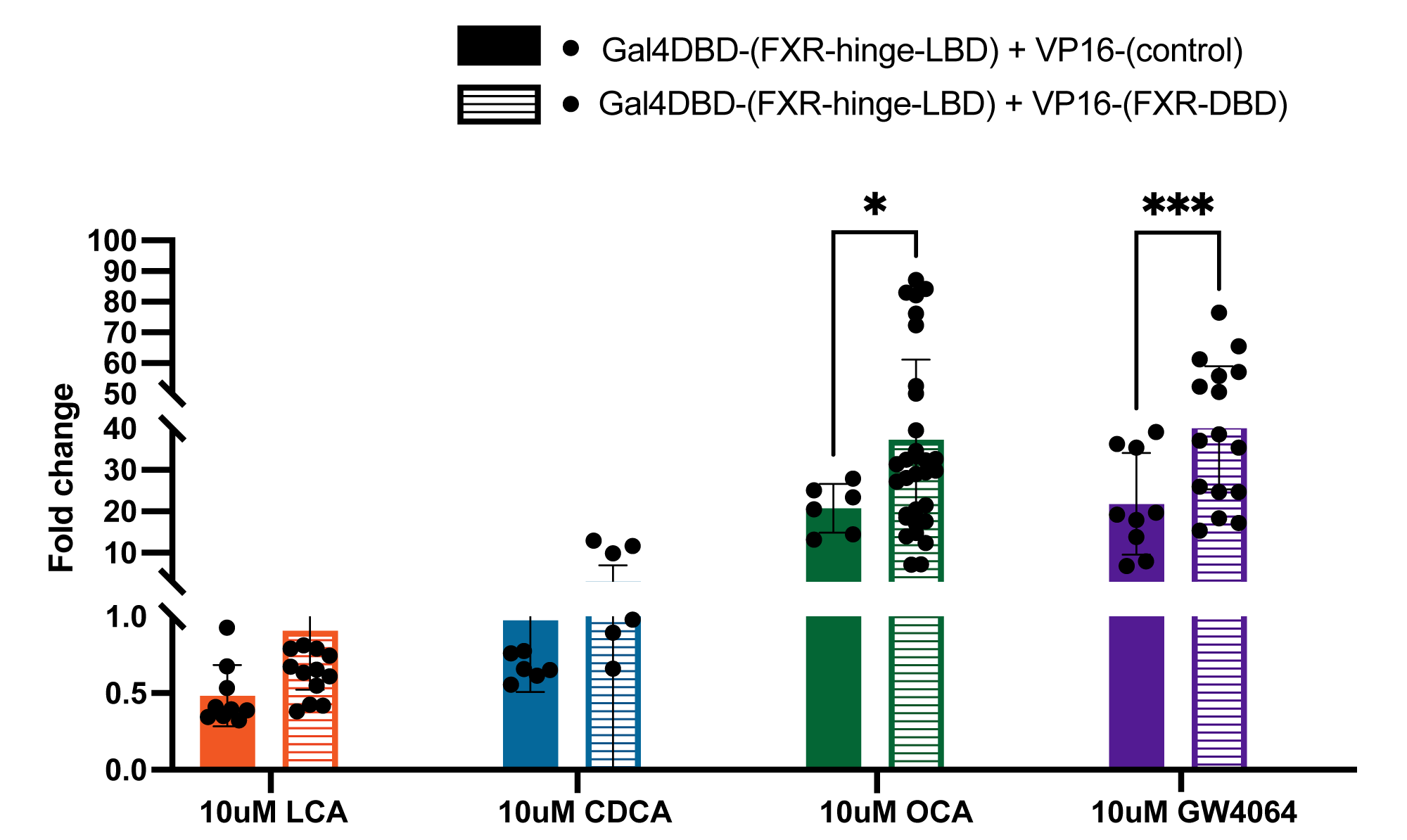

**Figure S7.** Mammalian two hybrid assay comparing protein-protein interactions in the presence and absence of the FXR-DBD. This experiment demonstrates that the increase in luciferase signal results from a hinge-induced interdomain interaction. Without the DBD, no increase in luciferase signal (over background) is observed.

**Table S2. Residues considered for calculating cluster parameters θ, r and d_v_ of different full-length crystal structures**

| Receptor(with PDB code) | Vector Origin  (residue) | Vector Termination (residue) | | Residues | | Rotational Angle θ  (degrees) | Interdomain distance r  (Å) | Vertical Displacement d_v_  (Å) |
| --- | --- | --- | --- | --- | --- | --- | --- | --- |
|  |  | V1 | V2 | DBD | LBD |  |  |  |
| PPARγ_3E00 | 415 | 228 | 422 | 107-175 | 208-477 | 69.98 | 38.166 | -3.92 |
| RARβ_5UAN | 350 | 177 | 357 | 80-150 | 176-408 | 76.99 | 35.056 | -4.59 |
| HNF4α_4IQR_2 | 305 | 141 | 312 | 49-120 | 141-368 | 81.39 | 67.286 | 34.59 |
| HNF4α_4IQR_1 | 305 | 143 | 312 | 49-124 | 143-368 | 97.32 | 38.429 | 14.34 |
| LXRβ_4NQA | 401 | 224 | 408 | 75-160 | 222-459 | 103.42 | 44.434 | 14.60 |
| RXRα_5UAN | 395 | 230 | 402 | 133-209 | 232-457 | 105.22 | 53.66 | 28.79 |
| RXRα_4NQA | 398 | 229 | 405 | 127-199 | 232-457 | 169.40 | 60.31 | 36.24 |
